# Connecting Chromatin Structures to Gene Regulation Using Dynamic Polymer Simulations

**DOI:** 10.1101/2023.11.07.566032

**Authors:** Yi Fu, Tianxiao Zhao, Finnegan Clark, Sofia Nomikou, Aristotelis Tsirigos, Timothée Lionnet

## Abstract

The transfer of regulatory information between distal loci on chromatin is thought to involve physical proximity, but key biophysical features of these contacts remain unclear. For instance, it is unknown how close and for how long two loci need to be in order to productively interact. The main challenge is that it is currently impossible to measure chromatin dynamics with high spatiotemporal resolution at scale. Polymer simulations provide an accessible and rigorous way to test biophysical models of chromatin regulation, yet there is a lack of simple and general methods for extracting the values of model parameters. Here we adapt the Nelder-Mead simplex optimization algorithm to select the best polymer model matching a given Hi-C dataset, using the *MYC* locus as an example. The model’s biophysical parameters predict a compartmental rearrangement of the *MYC* locus in leukemia, which we validate with single-cell measurements. Leveraging trajectories predicted by the model, we find that loci with similar Hi-C contact frequencies can exhibit widely different contact dynamics. Interestingly, the frequency of productive interactions between loci exhibits a non-linear relationship with their Hi-C contact frequency when we enforce a specific capture radius and contact duration. These observations are consistent with recent experimental observations and suggest that the dynamic ensemble of chromatin configurations, rather than average contact matrices, is required to fully predict productive long-range chromatin interactions.

## Introduction

The three-dimensional organization of the genome plays an important role in regulating gene expression by constraining long-range regulatory communication, for instance between promoters and enhancers (1). In proximity ligation-based methods such as Hi-C, two main genomic structures are apparent (2): Topologically Associating Domains (TADs), which are regions with high interaction frequency, and thus appear as squares around the diagonal in contact frequency matrices; and large A and B compartments, which are regions that preferentially interact with alternating regions but not neighbors, and thus appear as a checkerboard pattern in contact frequency matrices. On one hand, TADs constitute an emergent property of the chromatin fiber resulting from dynamic loop extrusion by Loop Extrusion Factors (LEFs), such as cohesin (3), and are bound by CTCF (CCCTC-binding factors) binding sites (4–6). On the other hand, compartments largely correlate with alternating heterochromatin and euchromatin domains (7).

Simulations combining polymer dynamics, loop extrusion by LEFs with fixed boundaries, and weak attractive domain interactions accurately describe genome organization, including compartments (3,8) and their dynamics (9,10). These simulations enable quantitative exploration of the mechanisms driving chromatin folding. Yet, given an experimental Hi-C dataset, there is a lack of simple and general methods to unbiasedly extract the parameter values of the corresponding polymer model. Recent work has compared experiments to a series of simulations based on a grid sweep of the parameters. Such a sweep provides an unbiased mapping of parameter space but is computationally costly (10). Others have used machine learning to find the parameters for the weak attractive domain interactions. However, determining the parameters for loop extrusion still requires systematically testing hundreds of parameter sets based on prior knowledge of CTCF binding (8).

Here, leveraging a polymer model that combines loop extrusion and domain interactions (3), we adapted the Nelder-Mead simplex optimization algorithm to build a method that can find the polymer parameters that best describe any input Hi-C data. We validated our strategy using *in silico* ground truth, published datasets featuring established biological perturbations, and DNA FISH data. Leveraging the temporal trajectories of chromatin generated by our simulations, we show that similar loci pairs with similar Hi-C contact frequencies can nonetheless exhibit dramatically different chromatin dynamics. Enforcing a specific capture radius and minimum duration for long-range communication, we show that long-range regulation varies non-linearly with Hi-C contact frequency across a range of plausible parameters.

## Materials and methods

### Cell Culture

Human peripheral blood CD4+ T cells were bought from Lonza (Lot No. 18TL156140) and thawed following the manufacturer’s protocol. The T cells were then cultured in RPMI media (Corning, #10-040-CV) supplemented with 15% fetal bovine serum (Invitrogen; 10082147), 1% penicillin/streptomycin (Thermo Fisher Scientific; 15-140-122), 1x glutamax (Fisher Scientific; 35-050-061), 1x non-essential amino acids (Fisher Scientific; 11-140-050), 1x sodium pyruvate (Thermo Scientific; 11360070), 1x beta-mercaptoethanol (Thermo Scientific; 21985023), and 30 U/ml IL2 (PeproTech; #212-12).

CUTLL1 cells are a gift from the Aifantis lab. The cell line was cultured in RPMI media supplemented with 10% fetal bovine serum and 1% penicillin and streptomycin.

### Imaging Sample Preparation

Cells were spun down to polylysine (Sigma-Aldrich; A-005-C) coated slides using a Thermo Scientific Cytospin 4 centrifuge at 1250 rpm for 5 min. Cells were then fixed in 4% paraformaldehyde (Electron Microscopy Sciences; #15713, diluted in PBS) at 4 °C overnight. After rinsing once with 1x PBS (Fisher Scientific 14-190-250), cells were permeabilized with 0.5% TritonX-100 (Bio-Rad #1610407, diluted in PBS) for 15 min at room temperature. Cells were rinsed again with PBS. The slides were used immediately for subsequent processes or stored in PBS at 4 °C overnight.

### DNA FISH

Custom DNA FISH probes were ordered from Arbor Biosciences (sequences in Supp. Table 1). To label DNA FISH probes, dual-HPLC purified, amino-modified oligos were obtained from IDT and labeled by incubation with NHS esters conjugated with Cy3 (Cytiva PA13101) or Cy5 (Cytiva PA15101) in 0.1 M sodium carbonate buffer (pH 9.0) overnight. Labeled oligos were purified with QIAquick nucleotide removal kit (Qiagen; #28304) and used as reverse transcription primers.

**Table 1.**
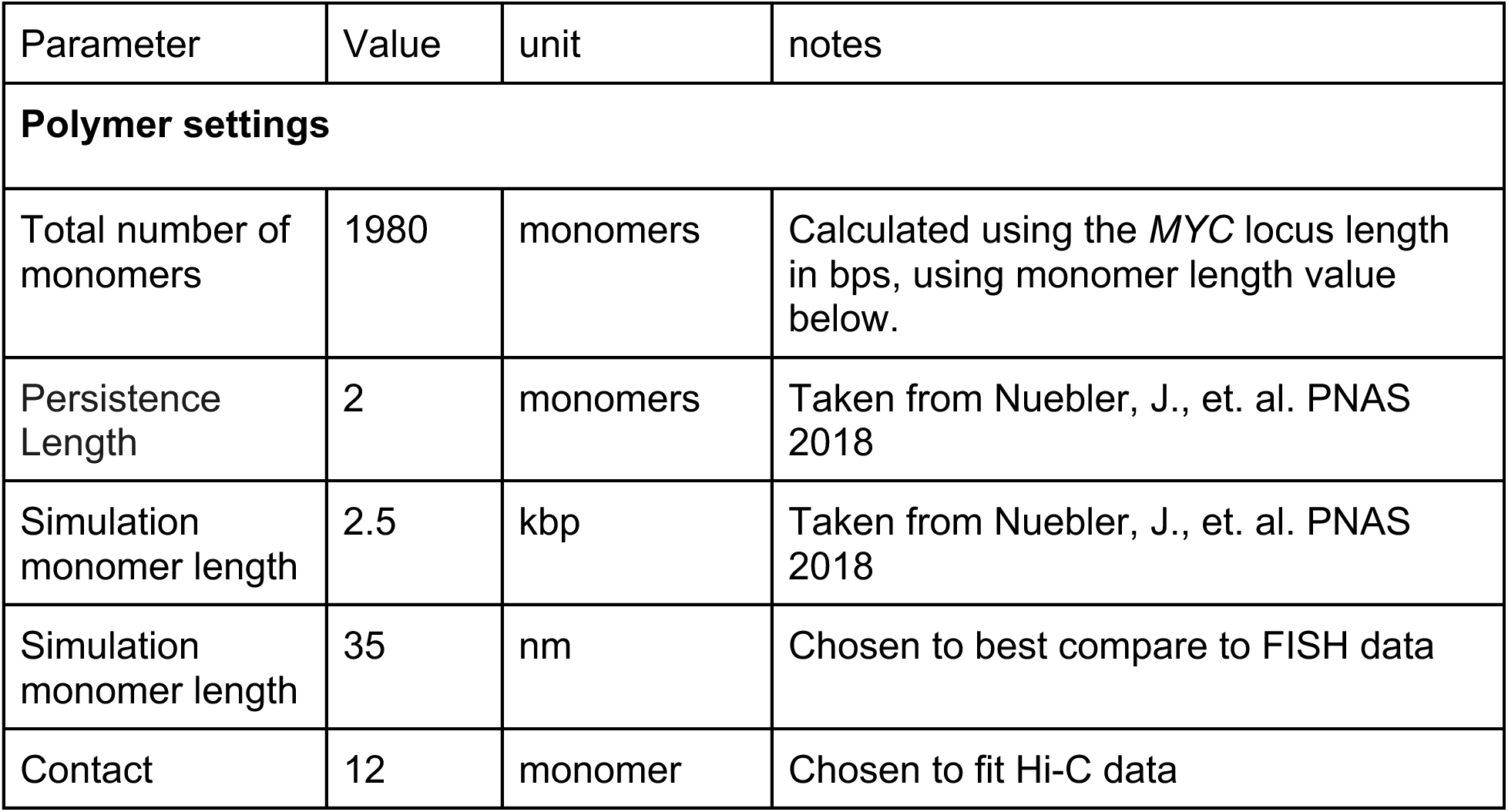
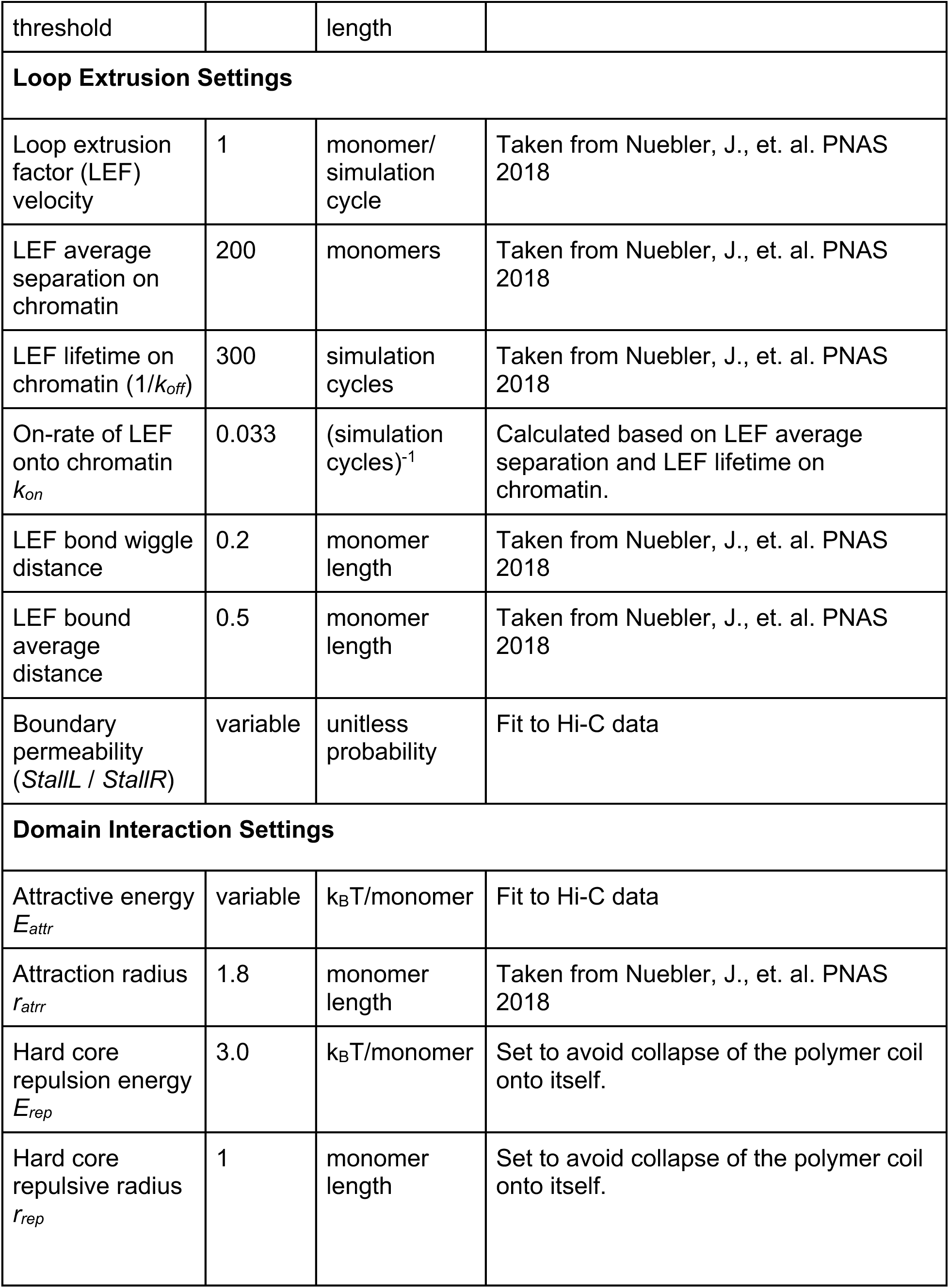
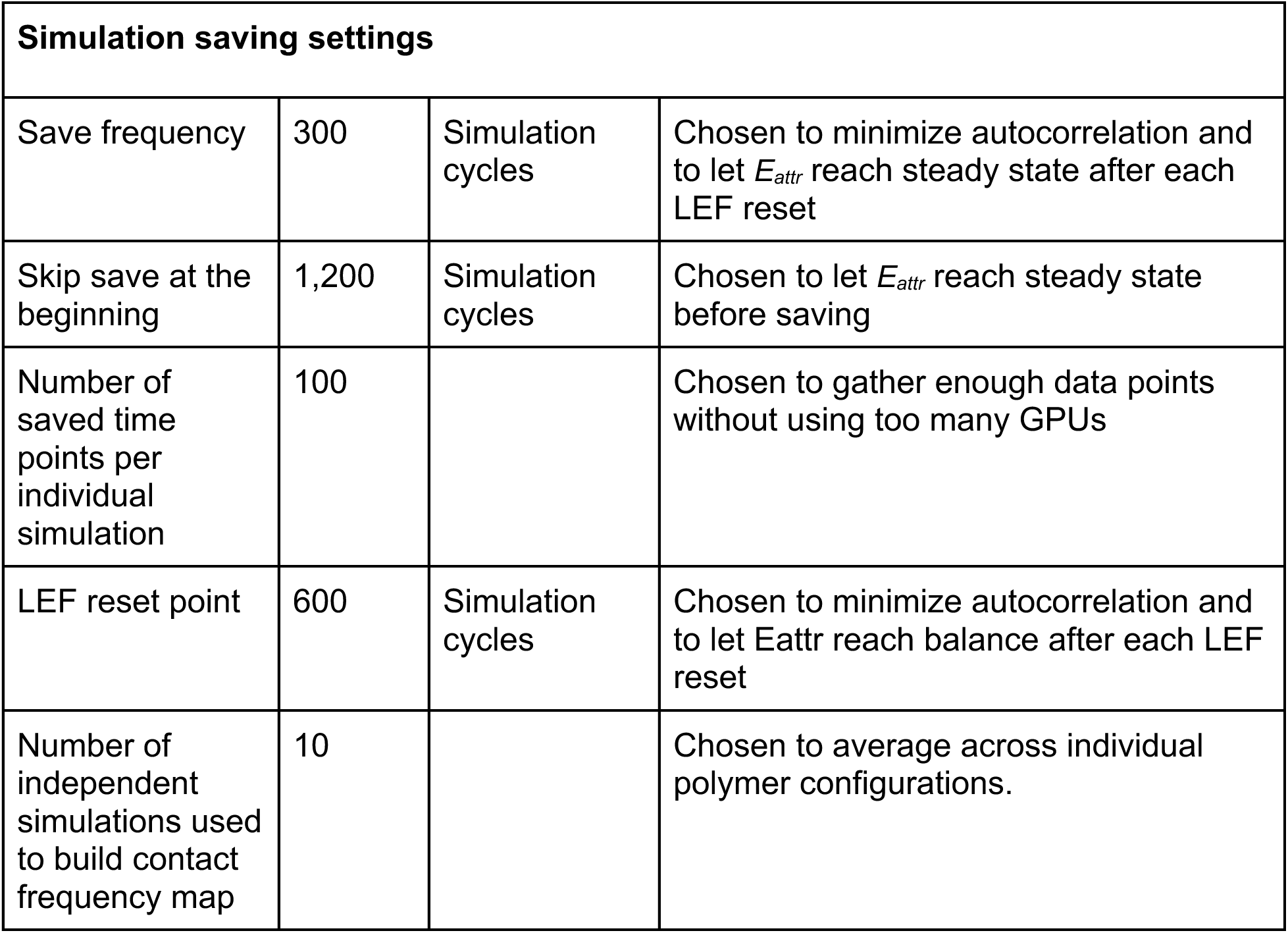
List of all simulation parameters and their values.

Probe labeling and DNA FISH were performed following the manufacturer’s protocol. The probes were resuspended in nuclease-free water at 0.07 ng/uL, and PCR amplified with KapaHiFi Hotstart Readymix (Fisher Scientific KK2601). Following PCR, a debubbling mix (10 uL KapaHiFi Hotstart Readymix, 1.2 uL myTag PCR primer mix, and 8.8 uL nuclease-free water) was added to the sample and incubated in a thermocycler with the following program: 1) 95 °C 3 min, 2) 98 °C 20 s, 3) 54 °C 15 s, 4) 72 °C 30 s, 5) repeat steps 2-4 one time. The product was purified with QIAquick PCR Purification Kit (Qiagen; #28104) and eluted in 30 uL of nuclease-free water.

To obtain single-stranded DNA FISH probes, after PCR purification, probes were *in vitro* transcribed with the MEGAshortscript T7 kit (ThermoFisher AM1354) for 4 h at 37 °C and purified with RNeasy Mini Kit (Qiagen; #74104). Following the in vitro transcription, the probes were reverse transcribed with Superscript IV Reverse Transcriptase (ThermoFisher 18090010). 52 ug *in vitro* transcription product and 2 nmoles labeled oligo were vacuum dried with Eppendorf Vacufuge Plus and resuspended in 44 uL nuclease-free water. The mixture was then combined with 15 uL 10 mM dNTPs (New England Biolabs; #N0447S) and 1 uL SUPERase-In (ThermoFisher AM2696) and incubated at 65 °C for 5 min. Then the mixture was combined with 6.5 uL nuclease-free water, 20 uL reverse transcription buffer, 10 ul 0.1 M DTT (Invitrogen; #1949355), 1 uL SUPERase-In, and 2.5 uL reverse transcriptase and incubated at 50 °C for 3 h. 11 uL Exonuclease I Buffer and 2 uL Exonuclease I (New England Biolabs; M0293S) was added following reverse transcription and incubated at 37 °C for 15 min. Then 12 uL 0.5 M EDTA (Invitrogen; #00486682) was added and incubated at 80 °C for 20 min, and the reverse transcription product was purified with Quick-RNA Miniprep Kit (Zymo; #R1054S). The probes were then digested with 4 uL RNaseH (New England Biolabs M0297S) and 4 uL RNaseA (Thermo Scientific; EN0531) in RNaseH buffer using the following program: 37 °C 2 h, 70 °C 20 min, 50 °C 1 h, 95 °C 5 min, ramp down to 50 °C at 0.1 °C/sec, 50 °C 1 h. Following digestion, the probes were purified again with Quick-RNA Miniprep Kit.

DNA FISH slides were incubated in 20% glycerol (Fisher Bioreagent; #172572, diluted in PBS) for 30 min, frozen in liquid nitrogen for 30 s, and thawed for 1 min at room temperature. The slides were again incubated in 20% glycerol for 20 min, frozen in liquid nitrogen for 30 s, and thawed for 1 min at room temperature. The slides were then incubated for 5 min in 0.1 N HCl (Fisher Scientific; S25354) and washed with 2xSSC (Sigma-Aldrich; 11666681001) 3 times, 1 min each. Then the slides were incubated for 30 min in hybridization buffer (30% formamide, Thermo Fisher Scientific; 15-515-026; 5xSSC, 9 mM Citric Acid, Sigma-Aldrich; 251275-500G and MKCF6319, pH 6.0; 0.1% Tween20; 1x Denhardt’s Solution; 40% Dextran Sulfate, Sigma-Aldrich; 42867-5G; 0.4 mg/mL BSA) at 37 °C. 10 pmol probes were diluted in hybridization buffer, incubated for 5 min at 70 °C, added to the slides, incubated at 85 °C for 5 min, and hybridized overnight in a 37 °C humidified chamber. The slides were then washed twice for 30 min with probe wash buffer (30% formamide, 5x SSC, 9 mM Citric Acid pH 6.0, 0.1% Tween20) at 37 °C and twice 5 min with 5x SSCT (SSC + 0.1% Tween20) at room temperature. Finally, the slides were rinsed with 0.5 ug/mL DAPI (Sigma-Aldrich; 32670-5MG-F) in 1xPBS, mounted in ProLong Gold (Thermo Fisher Scientific; P36930) and sealed with nail polish the next day (Electron Microscopy Sciences; #72180).

### Image Acquisition

Slides were imaged on a customized, MicroManager-controlled (11,12), epi-fluorescent Nikon microscope equipped with a Chameleon camera, using a 100x oil-immersion lens (NA=1.4). 10-15 image stacks were captured per sample with voxel size 73x73x250 nm. Slides were imaged using three channels: 405 (DAPI), 532 (Cy3), and 637 nm (Cy5) excitation lasers.

### Image Analysis

We first estimated the locations of the centroids of DNA FISH signals using AirLocalize(13), eliminating double detections. We then cropped out a small 3D region (20x20x10 pixels) around each approximate centroid, and subtracted the surrounding background intensity. We determined the centroid of each locus as the center of mass of the intensity distribution using the meshgrid function. We matched centroids from different channels if they shared the same nuclear mask (generated with CellProfiler (14)) and were mutual nearest neighbors, and calculated their mutual distance. Cells were categorized into G1 or G2 phase based on the number of FISH spots; one or two for G1, three or four for G2. Cells with no spots or more than four spots were excluded from the cell cycle analysis (statistics in Supp. Table 2). Matlab code (https://www.mathworks.com/) is available on GitHub: https://github.com/timotheelionnet/DNAFISH_analysis

### Polymer Simulations

Polymer simulations combining Langevin dynamics and loop extrusions were adapted from a previous publication(3). TAD boundary monomers were assigned two permeabilities, corresponding respectively to the probability of stopping a LEF moving forward (*stallL*) or backward (*stallR*; Fig. 1 C and Supp. Fig. 1 A). The interaction potential between monomers consists of a short-range repulsive hard core enforcing impenetrability, followed by an attractive well at intermediate distances. The well depth is set to the attractive energy *E_attr_*, a value specific to each TAD (Fig. 1 D and Supp. Fig. 1 A), and the attractive energy between each monomer pair is determined using the geometric mean of the attractive energies of both monomers. We set the well radius and hard core repulsion energy to values that recapitulate typical compartmental checkerboard patterns and kept them fixed throughout (Table 1).

**Figure 1.**
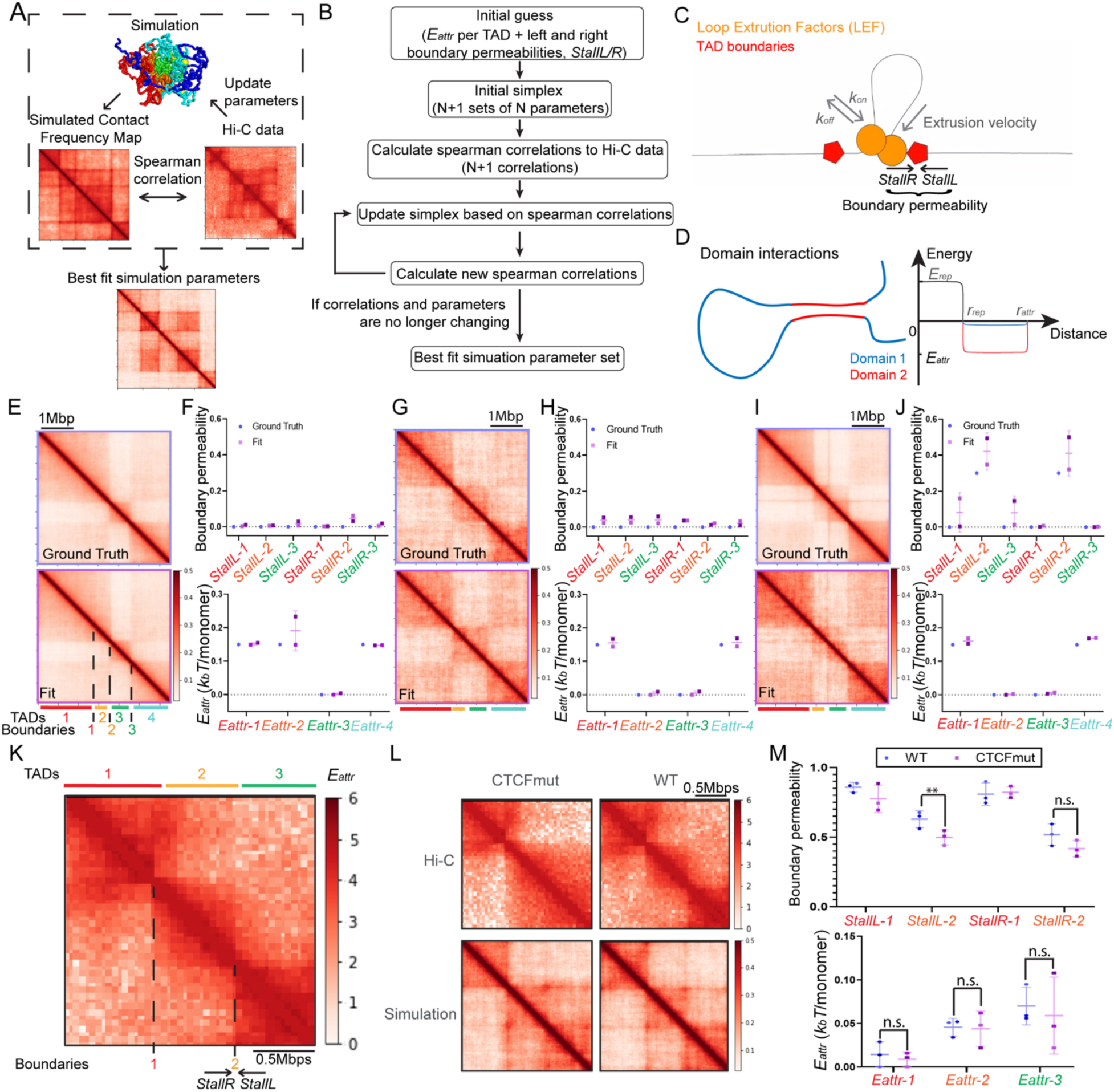
Simulation setup. A) Optimization principle. Contact frequency maps calculated from the simulated structures are compared to Hi-C data using Spearman Correlation; the simulations are then iteratively optimized to get the best-fit parameter set. B) Nelder-Mead algorithm flowchart. C and D) Loop extrusion and domain interaction models (parameters are listed in Table 1). E,G,I) Contact frequency maps of ground truth simulated data (top) compared to contact frequency maps of best-fit models (bottom). F,H,J) Comparison of the biophysical parameter values between ground truth and best-fit models. K) Model of a locus rearranged in CTCFmut (chr3: 59,800,000-61,840,000). Each TAD is assigned a specific *E_attr_* value. The intervening TAD boundaries are each assigned two directional boundary permeabilities (possibilities of stopping at the boundary in either direction, *stallL* and *stallR*). L) Top: Hi-C contact frequency maps in WT and CTCFmut. Bottom, best-fit simulations for WT and CTCFmut. M) Best-fit biophysical parameters of the simulations in L. **p < 0.05.

To optimize the simulation duration to streamline the parameter search (Supp. Fig. 1 B), we computed the autocorrelation function of the TAD2-TAD4 inter-TAD distance using the initial guess simulation parameters of the MYC locus in CUTLL. The simulation was saved every 5 simulation blocks. Based on the decay of the autocorrelation function of the simulated structures (Supp. Fig. 1 C), we saved structures every 300 blocks, and reset the positions of loop extrusion factors every 600 blocks; each block consists of a LEF position update (each LEF moves by one monomer or dissociates/reloads), followed by 250 Langevin dynamics iterations. Each simulation consisting of 10 parallel replicates takes <30 min on 10 GPUs.

To define the initial guess, we systematically adjusted the parameters one by one (*E_attr_* at 0.1 interval, *stallL*/*R* at 0.25 interval for each TAD/boundary) while keeping others at a fixed value. The best-fits from manual systematic adjusting were used as the initial guess for simplex.

The simplex leverages the Nelder-Mead method (https://docs.scipy.org/doc/scipy/reference/optimize.minimize-neldermead.html) as implemented in the scipy.optimize package (https://docs.scipy.org/doc/scipy/reference/optimize.html). To build contact frequency maps, we assigned contact between monomer pairs lying within a given distance threshold, set to 420 nm in best-fits (matching the lower resolution of HindIII Hi-C input data(15,16)), and 200 nm for the dynamic trajectories (matching the increased resolution of recent experiments using MboI capture-C(15,17)). These distance thresholds are within the range of similar threshold values used in previous publications comparing Hi-C with DNA FISH (Supp. Table 3).

We tested various methods to measure the similarity between simulated contact frequency maps and Hi-C maps (Supp. Fig. 2): Spearman correlation between simulated and Hi-C maps, Spearman correlation between 3-component PCA of simulated and Hi-C maps, squared distance between simulated Hi-C maps, and distance-corrected correlations between log-transformed simulated and Hi-C maps (8). All metrics converge to similar best fit parameters for the CUTLL Hi-C data. Since Spearman correlation between simulated and Hi-C maps required the least number of iterations, we used that method for all simulation fittings at the MYC locus. For smaller loci, distance-corrected correlations gave better agreement with the Hi-C data and were used instead.

The model used to fit into MYC Hi-C data consists of 1920 monomers representing chr8:126,720,000-131,680,000, with the TAD boundaries located at monomer 456 (chr8: 127,840,000 - 127,880,001), monomer 808 (chr8: 128,720,000 - 128,760,001), monomer 1178 (chr8: 130,160,000 - 130,200,001) and monomer 1592 (chr8: 130,680,000 - 130,720,001). The simulation code is available on Github: https://github.com/yf1037/ChromatinSimulation

### Dynamic simulation

To follow dynamics, we saved structure coordinates every five blocks. We converted simulation block units to minutes by comparing the temporal scaling of the mean squared displacement of monomers with published data (3).

### Hi-C Data Analysis

We downloaded CTCF mutant Hi-C data(18) from the GEO accession viewer (<u>GSM4041374</u>, <u>GSM4041375</u>), and processed them with Hi-C Bench(19) (https://github.com/NYU-BFX/hic-bench/) using default settings at 40 kb bin size. T-ALL(16) data were previously analyzed using Hi-C Bench(19). All coordinates are mapped to hg19.

### Significance tests

We estimated statistical significance between Cumulative Distribution Functions using two-sample, one-sided Kolmogorov-Smirnov tests. All other statistical tests used a two-tailed student t-test.

To test the sensitivity of Nelder-Mead simplex optimization, we estimated the standard deviation of optimizations based on 8 repeated optimizations of CTCFmut data (Supp. Fig. 1 E). We then set means of optimizations at a given difference, e.g. means at 0.4 and 0.6 for *ΔStallL* = 0.2, and simulated optimization results based on a normal distribution. Then, we calculated the probability of observing a significant difference using the simulated optimization results.

## Results

### A parameter optimization algorithm predicts the contributions of loop extrusion and domain interactions from Hi-C maps

The simulations leverage an established polymer model (3) combining the following ingredients: 1) Polymer dynamics based on a Rouse Model and Langevin dynamics; 2) Loop extrusion using possessive LEFs that are stochastically loaded onto the chromatin fiber (Fig. 1 C and Supp. Fig. 1 A) (TAD boundaries are set to discrete locations matching Hi-C TAD boundaries and assigned two directional permeabilities for LEFs: *StallL*,*StallR*); and 3) domain interactions via attractive interactions between monomers, modeled by a potential well whose depth (attraction energy, *E_attr_*) quantifies mutual affinity (Fig. 1 D and Supp. Fig. 1 A). To search for the best-fit parameter set for the loci of interest (*StallL*, *StallR, E_attr_* for each boundary/domain), we start the optimization with an initial guess set of biophysical parameters (see *Methods*) and conduct a series of polymer simulation replicates (Supp. Fig. 1 B). These simulations are combined to build a simulated Hi-C map in which pairs of monomers closer than a distance threshold are scored as making contact. We then measure the Spearman Correlation between the simulated Hi-C map and the experimental Hi-C map (Fig. 1 A). Using the Nelder-Mead simplex optimization algorithm (Fig. 1 B), we update the parameter values to find the parameter set maximizing the Spearman Correlation. We first validated the optimization method using ground truth maps built from simulation runs with known values of StallL, StallR, Eattr for each boundary/domain. After typically 70-90 iterations, the Nelder-Mead algorithm converges close to the expected parameters (Fig. 1 E-J).

Since CTCF binding blocks loop extrusion (18,20), simulations should predict a loss of boundary permeability when CTCF is perturbed. To validate whether simulations recapitulate known biology using experimental data, we leveraged recent Hi-C maps captured on a cell line homozygously expressing a CTCF mutant missing zinc fingers 9-11 (CTCFmut)(18,20). Cells expressing CTCFmut lose a subset of CTCF binding, resulting in chromatin rearrangements visible in Hi-C maps. We optimized parameters for one of the rearranged loci against Hi-C contact frequency maps measured in the WT and CTCFmut (chr3: 59,800,000-61,840,000) (Fig. 1 K). As expected, the simulation predicted a significant drop of 0.13 in boundary permeability in CTCFmut compared to WT (Fig. 1 L; Spearman Correlation: 0.85±0.02 for CTCFmut, 0.82±0.01 for WT), but no significant changes in *E_attr_* (Fig. 1 M). We repeated the optimization eight times (Supp. Fig. 1 D) to calibrate the sensitivity and specificity of the simulations against experimental data (Supp. Fig. 1 E).

Since the simulation contains stochastic components, we quantified the sensitivity to small changes (see *Methods*). With three independent optimization runs, there is a 70% chance of finding a significant difference in boundary permeability *StallL* between two simulated maps if the change is 0.2 or greater (Supp. Fig. 1 F). For *E_attr_*, the chance of finding a significant difference with a change of 0.1 *k_B_T* is 90% with three optimizations (Supp. Fig. 1 G). While the sensitivity further increases with the number of runs, the specificity is very high (low false positive rates) even with only two optimization runs.

To further test the generalizability of the simulation fitting, we also fitted the simulation to four other loci previously profiled by Hi-C (4). The simulation fittings recaptured the major features in all the loci (Supp. Fig. 3).

### The rearrangement of the *MYC* locus in leukemia cells is consistent with changes in *E_attr_*

Having established our parameter optimization approach, we went on to test whether it can formulate mechanistic predictions. We used the pathogenetic chromatin rearrangements at the *MYC* locus in leukemia as a model (16), and ran an optimization search for the biophysical parameters yielding best agreement between simulated and previously published Hi-C contact frequency maps of T-ALL (T cell acute lymphoblastic leukemia) patient samples, the model leukemia cell line CUTLL1 (Columbia University T cell Lymphoblastic Lymphoma 1), and naive T cells from healthy donors (Fig. 2. A).

**Figure 2.**
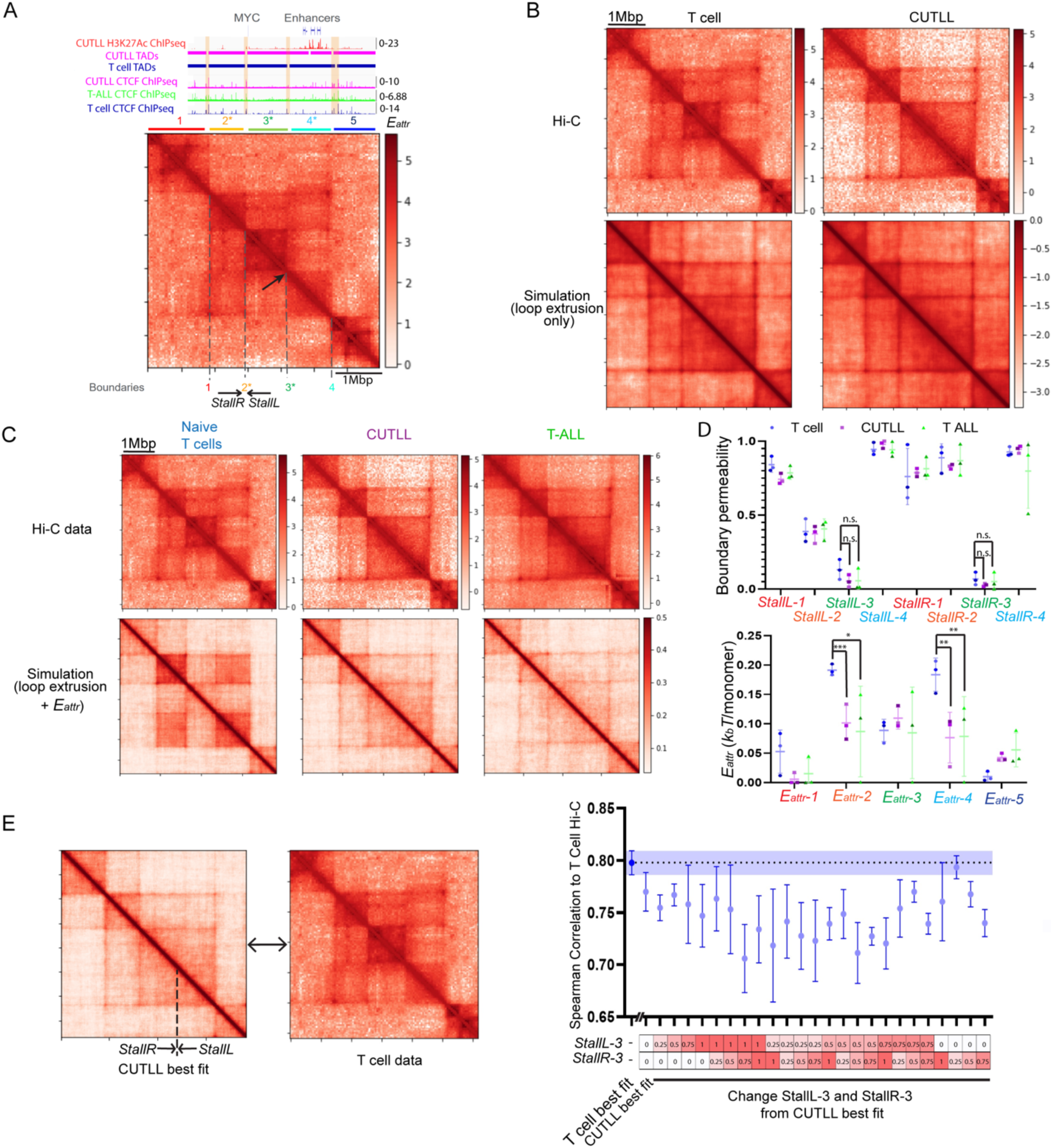
Simulation of the rearrangement of the *MYC* locus in T-ALL. A) Locus Model: five TADs surrounding the MYC gene are assigned distinct *E_attr_1-5,* while the four intervening boundaries are characterized by directional permeabilities (*StallL1-4, StallR1-4*). Boundary 3 (arrow) is disrupted in CUTLL1 and T-ALL patients. ChIP-seq and Hi-C map are plotted from a previous publication(16) (chr8:126,720,000-131,680,000). The log value of the contact matrix is plotted. B) Hi-C (top) and simulated (bottom) contact frequency maps of the MYC locus, using a polymer model featuring loop extrusion alone (no *E_attr_*). The log values of the contact matrices are plotted. C) Hi-C (top) and simulated (bottom) contact frequency maps of the MYC locus using the combined model with both loop extrusion and *E_attr_*. D) Best-fit parameter sets of the simulations in C. Left, TAD boundary permeability; Right, *E_attr_*. The result shows significant differences in TAD2 and TAD4 *E_attr_*. E) Correlations between Hi-C data and simulated contact frequency maps with boundary permeability of the disrupted boundary varying from 0 to 1 in 0.25 steps while other parameters are set to the best-fit of CUTLL1. T cell best-fit with high *E_attr_* best correlates with the Hi-C data. *p<0.1, **p < 0.05, ***p < 0.01.

Loss of CTCF binding at the boundary of the *MYC*-containing TAD (TAD boundary 3) in leukemia samples (16) suggests that a disruption in loop extrusion boundary could explain the observed changes in chromatin organization. In order to investigate whether loop extrusion changes alone could explain chromatin rearrangements, we ran the optimization using loop extrusion biophysical parameters only while omitting domain interactions. We separately ran optimizations to find the parameter values best describing Hi-C data from CUTLL1 or naive T cells. To our surprise, both optimizations converged to the same set of biophysical parameters, failing to recapitulate the distinct organization of the locus in T cells (Fig. 2. B; Spearman Correlation: 0.79±0.01 for CUTLL1, 0.67±0.02 for T cells).

Since loop extrusion alone appeared insufficient to explain the difference between the *MYC* locus structures of T cells and CUTLL1, we turned to the full model which includes both loop extrusion and *E_attr_*. With this model, we were able to obtain distinct Hi-C maps for the two conditions, each in agreement with its experimental counterpart (Spearman Correlation: 0.87±0.05 for CUTLL1, 0.84±0.07 for T cells, 0.93±0.03 for T-ALL; Fig. 2. C), regardless of the values of the parameters chosen to initialize the simplex (Supp. Fig. 1 I). In naive T cells, the best-fit exhibits stronger *E_attr_* between the TADs upstream (TAD2) and downstream (TAD4) of the *MYC*-containing TAD, compared to the same parameters in CUTLL1 and T-ALL (Fig. 2 D, Supp. Fig. 1 H). In other words, TAD2 and TAD4 preferentially interact with one another, but are segregated from the *MYC*-containing TAD3 in T cells. Interestingly, TAD4 super-enhancers are active in CUTLL1; the *E_attr_* disruption in leukemia could result in a TAD4 more permissive to promoter contacts (16).

To further confirm that changes in *E_attr_* but not boundary permeability underlie *MYC* rewiring, we generated contact frequency maps using a series of simulations set to different boundary permeabilities but keeping *E_attr_* constant, either set to the weak value observed in CUTLL1 (Fig. 2 E) or the strong value observed in T cells (Supp. Fig. 1 J). None of these simulated maps correlate with the T cell data better than the one with high *E_attr_*. Moreover, those with low boundary permeabilities correlate better than those with high boundary permeabilities. Together, these results suggest that changes in loop extrusion alone can’t explain the differences between CUTLL1 and naive T cells.

### Single-cell chromatin structures are consistent with *E_attr_* disruption at the *MYC* locus in leukemia

While contact frequency maps provide an informative readout of the average 3D organization of a locus, they are blind to the rich ensemble of conformations explored by chromatin (21). Polymer simulations, on the other hand, do sample the conformation ensemble: they not only predict the average organization of a locus, but also its variability across time and replicates, which should reflect the heterogeneity observed across cells. Having validated that the simulated *MYC* locus provides a realistic model of the average locus organization, we went on to test whether the ensemble of simulated structures recapitulates the heterogeneity in 3D organization measured in single cells. Simulations of the full model at the *MYC* locus predict that TAD2 and TAD4 are further apart in CUTLL1 than in T cells. In contrast, simulations of T cells where we only allow a gain of a loop extrusion boundary relative to the CUTLL1 best-fit (i.e. enforcing that *E_attr_* remains constant) do not lead to a substantial change in the inter-TAD distance (Fig. 3 A). We performed DNA FISH using probes tiling the two TADs in distinct colors and measured the distances between each TAD centroid in either cell type (Fig. 3 B, C, Supp. Fig. 4 A-C), confirming the simulation predictions that TAD2 and TAD4 are on average further apart in CUTLL1 compared to T cells (p = 4×10^-12^, two-sample one-sided Kolmogorov Smirnov test). This agreement suggests that the differences between CUTLL1 and naive T cells are caused by *E_attr_* changes but not loop extrusion. Importantly, not just the average distances, but the shape of the distance distribution across individual cells closely matches the predictions of the simulations in both cell types, further confirming that the simulations can predict heterogeneity across cells.

**Figure 3.**
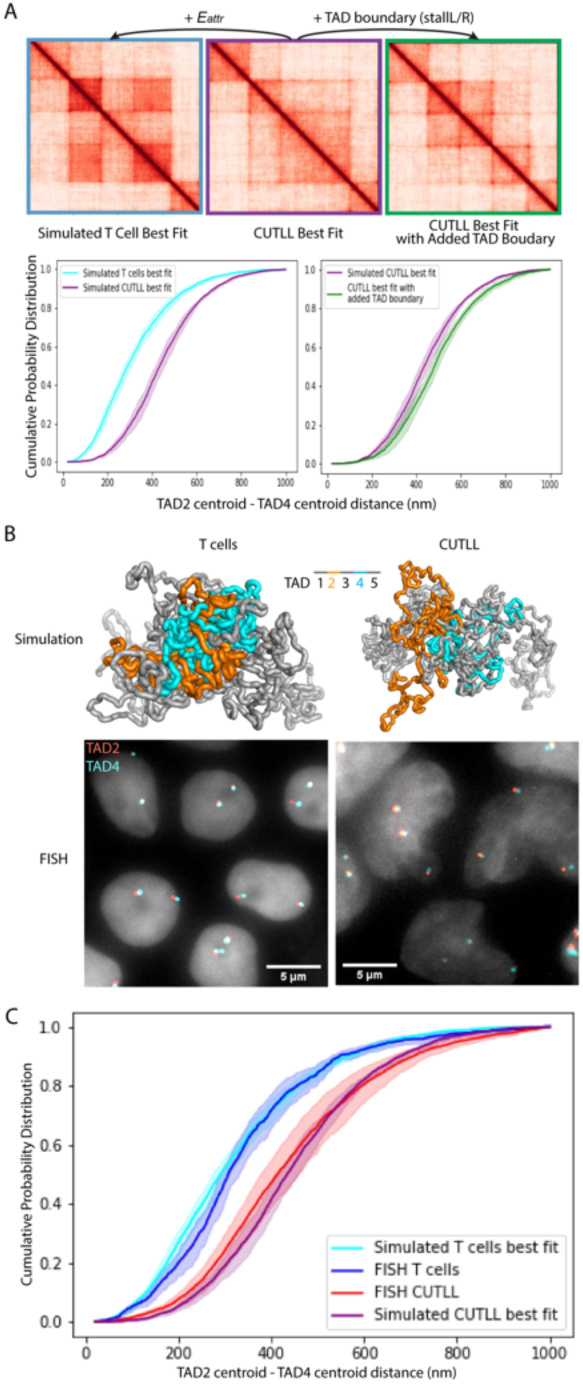
Simulations recapitulate the *MYC* locus disruption measured by DNA FISH. A) Top: Simulated Hi-C data; bottom: Simulated DNA FISH results; Center: CUTLL1; Left: simulated T cell with additional *E_attr_* compared to CUTLL1; Right: simulated data with an additional loop extrusion boundary compared to CUTLL1. B) Top: Representative snapshots of simulated *MYC* locus structures of T cells (left) and CUTLL1 (right) with TAD2 and TAD4 marked in orange and cyan respectively; bottom: representative FISH images of T cells (left) and CUTLL1 (right) using probes tiling TAD2 and TAD4 respectively. C) Cumulative probability distributions of the distances between the centroids of TAD2 and TAD4 in CUTLL1 and naive T cells obtained in simulations and in DNA FISH experiments. Shading represents standard deviations of each bin in the cumulative probability distributions (n = 3).

The faster cell cycle of CUTTL cells might constitute a possible confounder if the *MYC* locus confirmation is strongly cell cycle-dependent. To eliminate this possibility, we assigned individual CUTLL1 cells to distinct cell cycle stages according to the number of visible DNA FISH spots per nucleus (see *Methods*). The interTAD distances in G1 and G2 stages exhibit similar cumulative probability distributions (Supp. Fig. 4. D), suggesting that changes in *E_attr_* rather than differences in cell cycle phases explain the *MYC* rearrangement in leukemia cells.

### Pairs of loci with the same contact probability exhibit dramatically different looping dynamics

Because polymer simulations recapitulate complex chromatin organization features, from sensitivity to key perturbations (3) to single-cell distributions (8) and dynamic chromatin behavior (10,22), we reasoned that they might help investigate how the chromatin dynamic ensemble relates to average contact frequency maps, a widely used observable. Using the best-fit to the *MYC* locus in CUTLL1 as a testing ground, we simulated chromatin temporal trajectories to characterize how dynamics shape the interactions of the *MYC* promoter with its surroundings (Fig. 4 A and B). While the *MYC* locus constitutes a realistic chromatin landscape within which we explore general principles driving long-range interactions, we stress that we do not intend here to recapitulate the specifics of *MYC* expression regulation.

**Figure 4.**
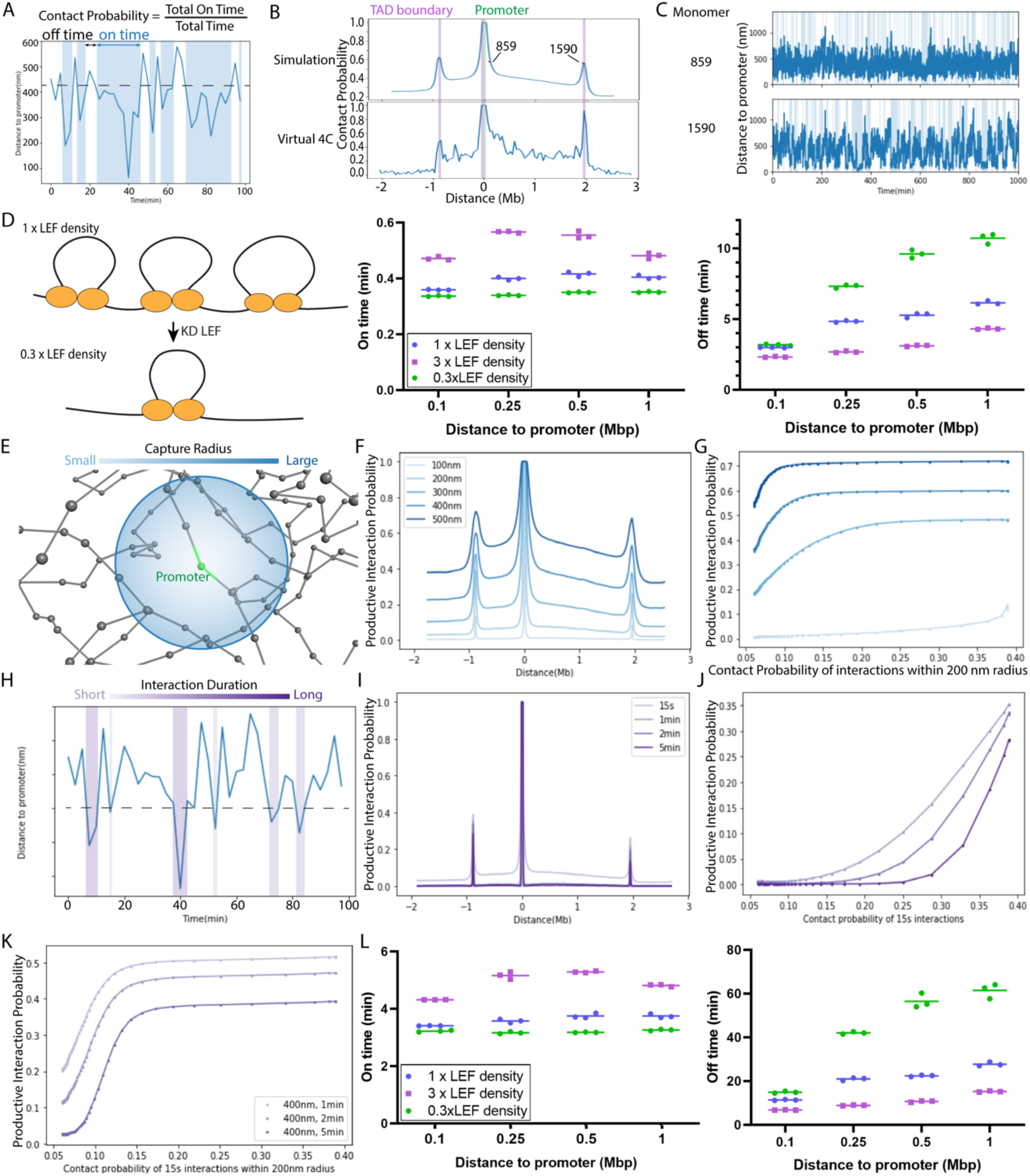
Simulated chromatin dynamics. A) Contact frequency definition. On-times are defined as time intervals during which two loci are closer than the set capture radius. B) Frequency of contact (<400 nm) with the promoter (monomer 811, green) as a function of genomic position across the locus in simulated data (top) and CUTLL1 Hi-C data (bottom). Two loci of interest (monomers 859, 1590) exhibit similar contact frequency despite very different distances to the promoter. C) Example simulated time traces of the distance between the promoter and either locus marked in B. D) Left: LEF knock-down/overexpression is modeled by enforcing different LEF densities in the simulation. Center, Right: Average On- and Off-time durations of contacts (<200 nm) between the promoter and monomers separated by different genomic distances, for different LEF densities. Each data point is the average of 10 independent simulations. E) Distance-gated productive interactions occur when a locus is within a set capture radius from the promoter. F) Simulated frequency of productive interactions with the promoter (located at the distance origin) as a function of genomic position across the locus, for different capture radii. G) Frequency of productive interactions as a function of the simulated Hi-C contact frequency, for different capture radii (Hi-C capture radius: 200 nm; minimum duration set to 15s for all curves; color scheme matches that of F, from light to dark: 100 nm, 300 nm, 400 nm, 500 nm) H) Duration-gated productive interactions occur when a locus is within a set capture radius from the promoter for at least a minimum duration. I) Simulated frequency of productive interactions with the promoter (located at the distance origin) as a function of genomic position across the locus, for different minimum durations. J) Frequency of productive interactions as a function of the simulated Hi-C contact frequency, for different minimum durations (Hi-C minimum duration: 15 s; contact radius set to 200 nm for all curves; color scheme matches that of I, from light to dark: 1 min, 2 min, 5 min). K) Frequency of productive interactions as a function of Hi-C contact frequency when enforcing time- and distance-gating; (productive interaction capture radius and minimum duration for each curve are indicated in the box; Hi-C capture radius and minimum duration: 200 nm, 15 s). L) Average On- and Off-time durations of productive interactions (capture radius: 400 nm; minimum duration: 2 min) between the promoter and monomers separated by different genomic distances, for different LEF densities. Each data point is the average of 10 independent simulations.

We first selected two loci with similar contact frequencies to the *MYC* promoter in simulated Hi-C data (monomer 859 and 1590, respectively located 120 kbp and 1.9 Mbp downstream of the *MYC* promoter at monomer 811; Fig. 4 B). Surprisingly, simulated contacts (<200 nm) between the promoter and either locus exhibited very distinct dynamics: contacts with monomer 859 are short and frequent, while contacts with monomer 1590 are long but infrequent (Fig. 4 C and Supp. Fig. 5 A).

We hypothesized that the long infrequent promoter contacts of monomer 1590 are driven by loop extrusion, while the frequent short promoter contacts of monomer 895 are mediated by the thermal motion of the chromatin fiber. Indeed, decreasing the density of LEFs on chromatin threefold results in minimal effects on the dynamics of promoter contacts with close monomers, but significantly impacts the dynamics of promoter contacts with far away monomers (Fig. 4 D and Supp. Fig. 5 B). We also tested perturbing the lifetime of LEFs on chromatin which again impacted the contacts with monomers far away from the promoter but not those with closeby monomers (Supp. Fig. 5 B). These findings illustrate that Hi-C contact frequency has limited power to predict looping dynamics.

### Time- and distance-gating regulate chromatin contacts non-linearly

The transcription output of a reporter gene exhibits a sigmoidal response to the frequency of contacts between its promoter and a distal enhancer, as measured by Hi-C (17,23), although the molecular mediators of this relationship remain unknown. Cross-linking used in Hi-C likely captures transient contacts because DNA binders that reside on chromatin for ∼5-100 seconds (24) are efficiently captured by crosslinking in chromatin immunoprecipitation techniques. Comparisons between Hi-C and DNA FISH suggest that loci crosslink in Hi-C if they lie within ∼100-400 nm from each other, with different variants of the Hi-C protocol providing slightly different spatial resolutions (Supplementary Table 3)(15,25). Yet, it remains unclear how close two loci need to come together in order to transfer regulatory information, and how long the contacts need to last in order to yield productive activation of a target gene. Based on the complex dynamics of our model locus, we reasoned that non-linearities in chromatin contact kinetics might account for the sigmoidal response of transcription with Hi-C contacts observed experimentally.

While Hi-C captures transient and close contacts (15s, 200 nm), the capture radius and minimum duration of productive interactions between distal chromatin loci remain undetermined. We first tested how the frequency of productive interactions between loci would change as a function of the capture radius (Fig. 4 E) or the minimum duration (Fig. 4 H). The frequency of productive interactions with the promoter as a function of genomic distance varied non-linearly with the capture radius or the minimum duration (Fig. 4 F, I). Interestingly, the frequency of productive interactions also displayed a non-linear relationship with the frequency of transient, close contacts mimicking Hi-C capture (15s, 200 nm) for a series of capture radii and minimal durations (Fig. 4 G, J, Supp. Fig. 5 C). We also confirmed that, independent of gating in radius and duration, loop extrusion drives long, infrequent interactions between distant loci, but not short frequent interactions between nearby loci (Supp. Fig. 5 D and E).

When combining changes in minimum duration and capture radius, we found that the frequency of long interactions within a large capture radius (> 1 min, 400 nm) exhibited a sigmoidal response to the frequency of close contacts mimicking Hi-C capture (15s, 200 nm; Fig. 4 K and Supp. Fig. 5 F and G). Recent observations show that cohesin depletion results in a sharp drop in the frequency of productive bursts of transcription, with little change in their duration or amplitude (23). Simulated productive interactions (400 nm, 2 min) echoed these findings, predicting less frequent contacts, but no major changes in their duration in the presence of LEF depletion (Fig. 4 L). Interestingly, the predicted effect of cohesin depletion on the frequency of productive interactions is much more pronounced at large genomic distances, consistent with findings that loop extrusion is crucial for enhancer function only at large distances (26,27).

The dependence of contact dynamics on loop extrusion in our simulations of *MYC* differs from that previously observed for two TAD boundaries (45). To check whether the different results are the product of different simulation models, we simulated contact dynamics across two TAD boundaries matching the locus of (45). Our simulations recapitulate the distance distribution and loop extrusion dependence previously observed (Supp. Fig. 6), establishing that the differences between the two systems are biological. Thus, while loop extrusion controls both the frequency and duration of contacts at TAD boundaries, it exerts a more nuanced effect on the frequency of contacts in loci pairs like the *MYC* locus that might better reflect typical enhancer-promoter pairs.

## Discussion

In this study, we developed a parameter optimization workflow to predict the respective contributions of loop extrusion and *E_attr_* from Hi-C maps. We were able to fit biophysical models of the chromatin fiber to Hi-C data using the Nelder-Mead algorithm, confirming that a combination of the two processes provides a tractable but realistic depiction of the large-scale features of chromatin folding (3,8). Importantly, best-fit simulations predict not only population-averaged observables such as Hi-C contact frequency but also single-cell variability, measured with DNA FISH.

Besides the modeling proposed here, the String Binders Switch (SBS) model (8) of chromatin has recently been used to fit experimental 3D organization data. In this model, different segments of chromatin are assigned a different binder type and can phase-separate into different globular compartments, compartmental TADs, or sub-TAD structures. The SBS approach successfully models chromatin organization in detail, at the cost of a larger, more complex set of parameters. The parameter optimization is computationally costly, requiring several days of computational time for 13 parameters used in our *MYC* locus model. Increasing the number of parameters would likely increase optimization time and/or require additional constraints, such as prior knowledge of CTCF binding (8). The model presented here offers the advantage of simplicity and only needs Hi-C data as input for parameter optimization. Future work extending this framework to single cell readouts out chromatin architecture (e.g. single-cell Hi-C or chromatin tracing) holds promise to further constrain chromatin models.

Validating the model on the oncogenic rewiring of the *MYC* locus, we found that the rewiring is consistent with a loss of *E_attr_* rather than a loss of loop extrusion permeability (Fig. 2 and 3). While major changes in loop extrusion permeability are ruled out by the simulations, we cannot rule out small changes in loop extrusion permeability that might fall under the method’s detection threshold. Such small changes are unlikely to explain the observed rewiring (Fig. 2 E, Supp. Fig. 1 J). In addition to blocking loop extrusion by cohesin, CTCF facilitates interactions of different compartments (28). It is thus possible that CTCF might play this non-canonical role instead of TAD insolation when bound to the boundary between TAD3 and TAD4 in T cells.

One possible explanation for the disruption of *E_attr_* at the MYC locus could be the loss of histone methylation marks. Polycomb Repressive Complex 2 (PRC2) establishes initial H3K27me2/3 and spreads H3K27me3 on chromatin (29). Polycomb Repressive Complex 1 (PRC1) reads these H3K27me3 marks and induces the formation of chromatin compartments by forming oligomers (29). PRC2 is frequently mutated or silenced in T-ALL, and is inactive in CUTLL1 (30). Consistently, T-ALL and CUTLL1 exhibit low H3K27me3 levels genome-wide (30), which could lead to the *E_attr_* changes observed at the *MYC* locus.

Analyzing the dynamics of chromatin structures, we observed that pairs of loci with similar Hi-C contact frequencies exhibit distinct looping dynamics: nearby loci pairs are dominated by thermal fluctuations while loci far apart exhibit more infrequent interactions mediated by loop extrusion. These observations raise the question of how the temporal dimension - largely missed in Hi-C - factors into long-range regulation. Transcription activation from a distal enhancer exhibits a sigmoidal relationship to the Hi-C contact frequency with its target promoter (17). The simulations presented here suggest that the sigmoidal relationship could emerge from productive interactions being limited to long events (> 1min) within a larger capture radius (∼400 nm) than that of Hi-C. In support of this idea, the time- and distance-gated model proposed here could recapitulate several observations: the increased dependence of loop extrusion for transcription activation from long distances (27,32); the stronger effect of loop extrusion perturbations on the frequency of activation intervals compared to their duration (23); the fact that Sox2 transcription exhibits little temporal correlation with distance to its SCR enhancer - since the SCR lies within <400 nm of the promoter the vast majority of the time, the model predicts near uninterrupted enhancer-promoter communication (33,34); the strong temporal correlation between enhancer-promoter distance and transcription activity for artificial systems where the enhancer regularly explores distances outside of the ∼400 nm capture radius from the promoter (35,36). While other mechanisms have also been proposed to account for the non-linearity observed between chromatin contact matrices and transcription regulation (17,37,38), the simulations presented here demonstrate that the choice of a specific contact observable as a metric is not a neutral decision, but in itself can introduce non-linearities. More importantly, these findings put into question the use of a single metric (e.g. Hi-C contact) to describe a complex dynamic ensemble (the chromatin fiber). A unidimensional metric has obvious practical advantages, yet it remains unknown which features of chromatin’s dynamic ensemble are interpreted by the cell to orchestrate long-distance regulation, and whether those features are captured by Hi-C contacts linearly and unambiguously.

While the values of the time and distance gates presented here are by necessity approximative due to the simplicity of the model, they represent plausible estimates. The contact radius of ∼400 nm, sufficient to generate a sigmoid relationship with Hi-C contacts in simulations, is tantalizingly close to the typical size of transcription clusters or hubs: dynamic, non-stoichiometric assemblies of transcription factors and co-activators observed around active genes and enhancers (34). Interestingly, the 400 nm capture radius also coincides with the typical distance under which promoters are more likely to burst in sync (39). These findings are consistent with the idea that regulatory information transfer might not rely on direct molecular contacts between chromatin segments, but rather on the continued presence of distal elements within a relatively large radius. Transcription regulators might then diffuse at a very short range between chromatin segments, possibly confined within local nano environments characterized by specific composition and biochemical properties (40). In such a model, why would only interactions longer than ∼2 minutes become productive? One possibility is that time-gating ensures regulatory specificity: increased dwell times of transcription activators at regulatory targets are stronger predictors of transcription output than increased average occupancy (41), suggesting the presence of kinetic proofreading steps in the transcription cycle (42–44). The minimal duration of ∼2 min predicted by the model allows a few typical transcription factor binding events to take place, which is likely sufficient to enable proofreading of the transcription binding events.

Our parameter optimization can be adapted to build biophysical models of any locus of interest. Despite the model simplicity, the best-fit simulations are sufficient to predict the contribution of loop extrusion and domain interactions, as well as single-cell variability from Hi-C data. Modeling dynamics enables testing mechanistic relationships between chromatin dynamics and transcription regulation. As more experimental results emerge to define simulation parameters, updates to the model should further increase its power.

## Supporting information

FISH probe sequences

## Acknowledgements

We would like to thank the Aifantis lab for sharing reagents and useful discussions, and the Applied Bioinformatics Laboratories (ABL) for providing bioinformatics support and helping with the analysis of the CTCFmut Hi-C data. ABL is a shared resource partially supported by the Cancer Center Support Grant P30CA016087 at the Laura and Isaac Perlmutter Cancer Center.

The computational requirements for this work were supported in part by the NYU Langone High Performance Computing (HPC) Core’s resources and personnel, and in part through the NYU IT High Performance Computing resources, services, and staff expertise. We thank the Fenyo lab for their assistance with HPC setup.

## Author Contributions

Y.F. and T.L. designed the experiments. F.C. developed the optimized DNA FISH protocol and performed some of the DNA FISH experiments. S.N. and A.T. analyzed H3K27me3 ChIP-seq data and some of the Hi-C data. Y.F. performed all other experiments and computations. T.L. supervised the findings of this work. All authors contributed to the final manuscript.

