## Supplementary figures for "Connecting Chromatin Structures to Gene Regulation Using Dynamic Polymer Simulations"

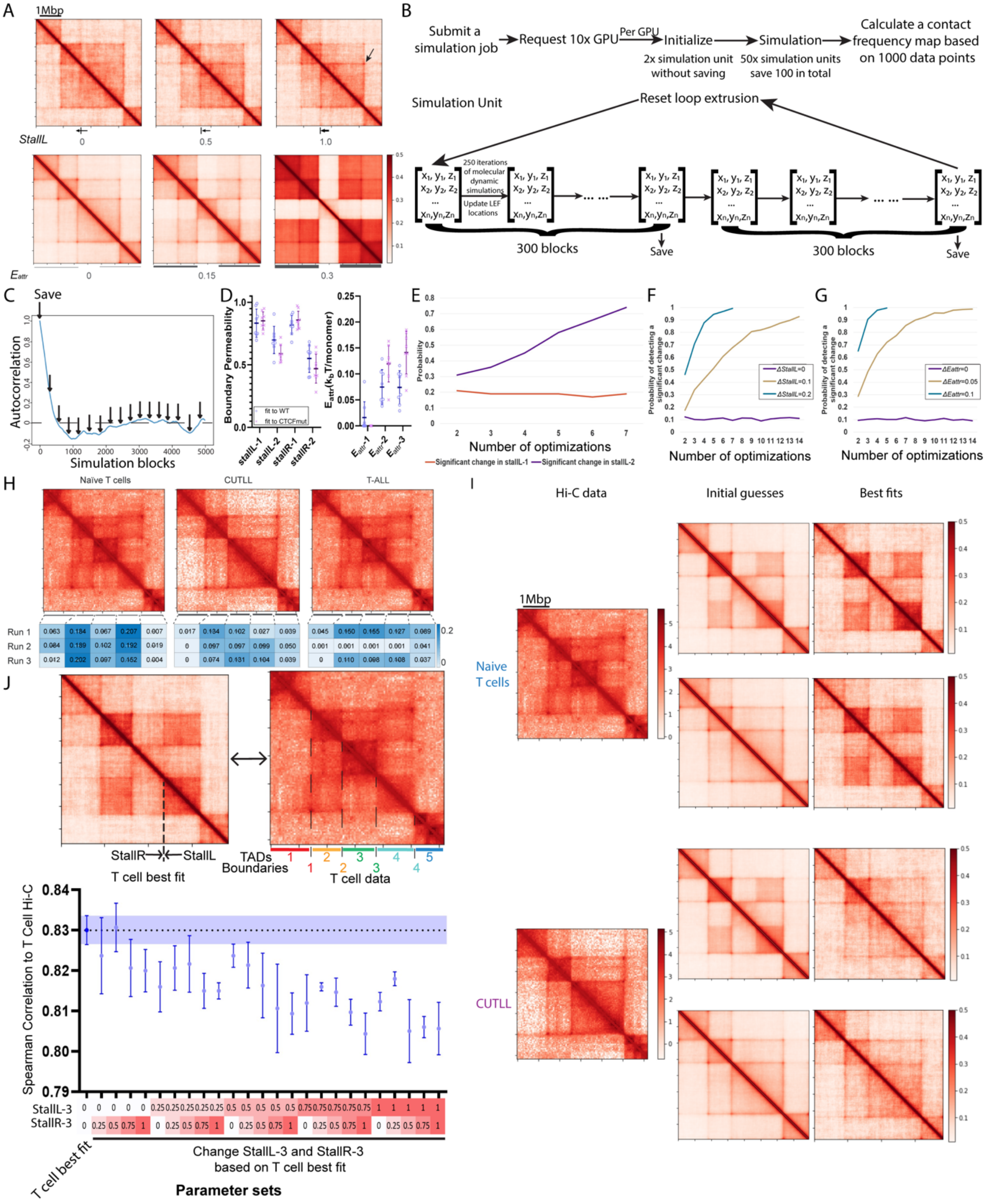


Supplementary Figure 1. Simulation validation. Related to Figure 1 and 2. A) Effects of changing loop extrusion boundary strength (StallL, top) and $E_{attr}$ (in $k_BT$ units, bottom). Parameter values increase from left to right. A strong boundary permeability leads to a stripe and a corner dot (arrow), while a strong $E_{attr}$ leads to a marked checkerboard pattern. B) Schematic of simulation blocks. 10 GPUs were used to simulate 50 simulation units and save 100 data points each. The simulated structures were saved every 300 blocks; the first two were skipped to allow the initial equilibration of the structure. C) Autocorrelation of simulated chromatin structures as a function of the temporal lag (in units of simulation blocks), in the absence of loop extrusion reset. The autocorrelation vanishes after ~600 blocks. D) Simulation reproducibility against the experimental data in Fig. 1L. Each data point represents the value of a parameter obtained in one round of optimization. E) Probability of detecting a significant true positive (significant change in *StallL-2*) or false positive (significant change in *StallL-1*) as a function of the number of optimization rounds for the data in Fig. 1L. Increasing the number of repeats increases the specificity while sensitivity is high even with two repeats only. F and G) Sensitivity and specificity of the model as a function of the magnitude of a change in either boundary permeability (Δ*StallL,* F) or $\Delta E_{attr}$ (in units of $k_B T$, G) (see *Methods*). H) $E_{attr}$ of the MYC TADs obtained in three separate optimization runs are shown as heat maps under the Hi-C matrices corresponding to each of the experimental conditions fitted. Despite differences in absolute $E_{attr}$ values between simulation runs, T cells reproducibly form a clear checkerboard pattern (*i.e.* TAD2 and TAD4 exhibit higher $E_{attr}$ than TAD3; left). The checkerboard pattern is never observed in CUTLL1 or T-ALL. I) Simulation optimization using different initial guesses gave similar results. Left: experimental Hi-C contact frequency maps used in simulations. Center: contact frequency maps calculated from structures simulated using the initial guess parameter values. Right: contact frequency maps calculated from structures simulated using the best-fit parameter values. J) Correlations between T cell Hi-C and simulated contact frequency maps of the MYC locus obtained by systematically varying the permeabilities of TAD boundary 3 (*StallL3, StallR3*), while keeping other parameters set to their values in the T cell best-fit. No increase in boundary strength results in a significantly better fit to the Hi-C data relative to the T cell best-fit.

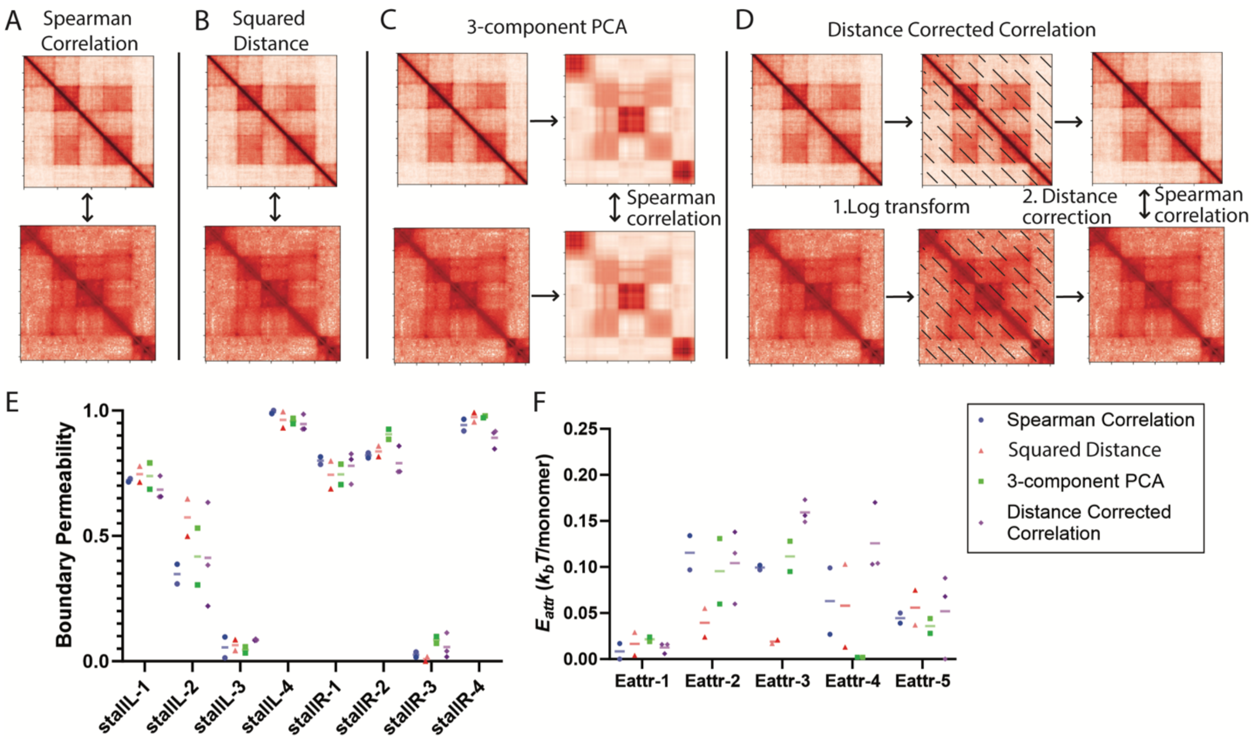


Supplementary Figure 2. Metrics comparing simulated and Hi-C data. A-D) Schematic of different metrics used to evaluate similarities between simulated and Hi-C data. A) Spearman correlation calculates Spearman correlation between simulated and Hi-C data directly. B) Squared distance calculates squared distance between simulated and Hi-C data directly. C) 3-component PCA takes the first 3 principal components from principal component analysis (PCA) of simulated and Hi-C data, and then calculates the Spearman correlation. D) To calculate distance corrected correlation, the simulated and Hi-C data are first log transformed, and then distance corrected by subtracting the average of each diagonal before calculating the Spearman correlation. E and F) CUTLL best fit boundary permeabilities (E) and attraction energy (F) using each of the metrics illustrated in (A-D). Different metrics converge to similar best fits.

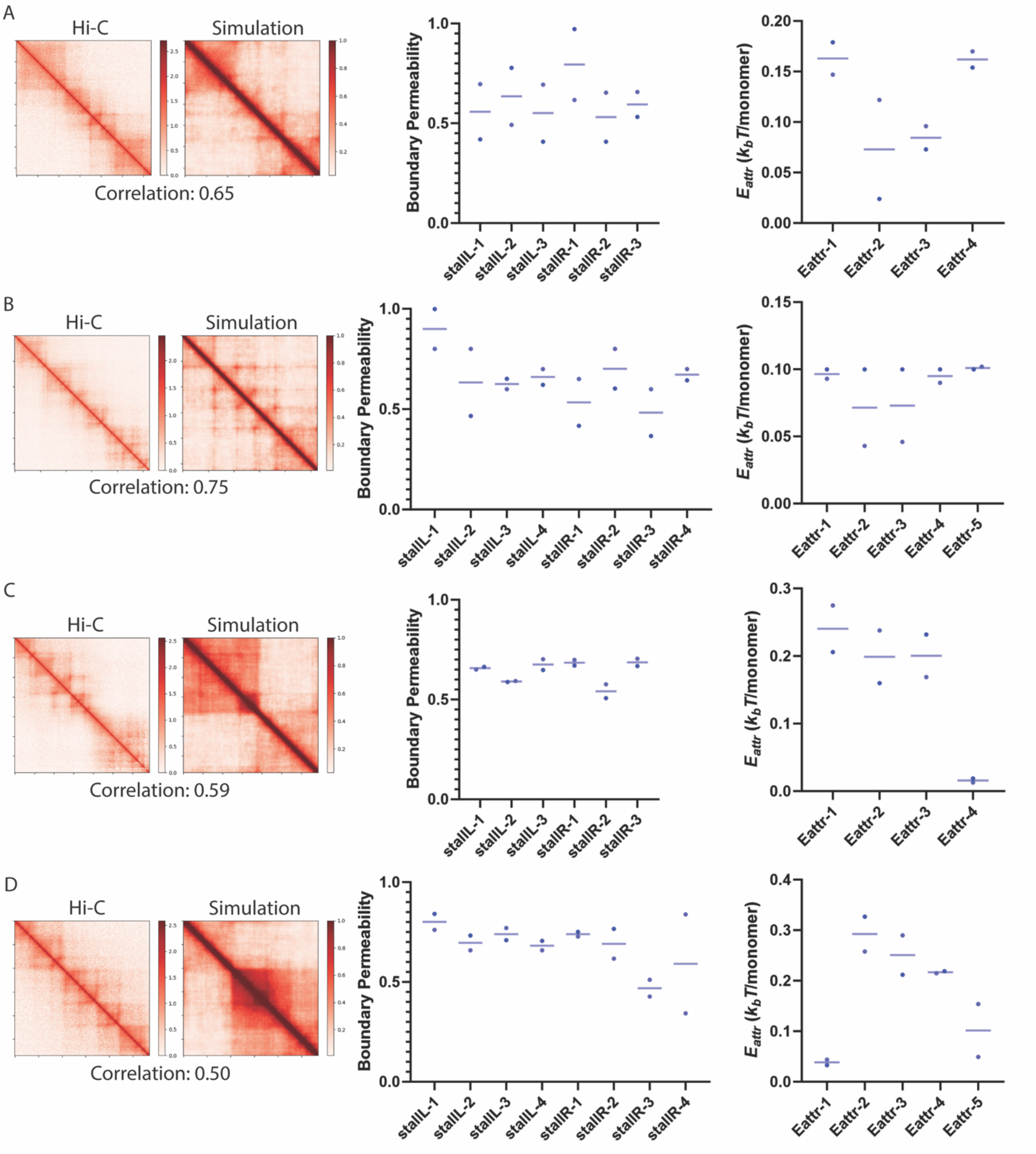


Supplementary Figure 3. Simulation fittings of additional loci using distance-corrected correlations. Left, Hi-C and simulated contact frequency maps of K562 cells from (4). Right, best-fit parameter sets. A) Chr19: 48,610,000 - 50,030,000. B) Chr3: 127,240,000 - 129,865,000. C) Chr12: 53,890,000 - 55,605,000. D) Chr19: 12,200,000 - 13,790,000.

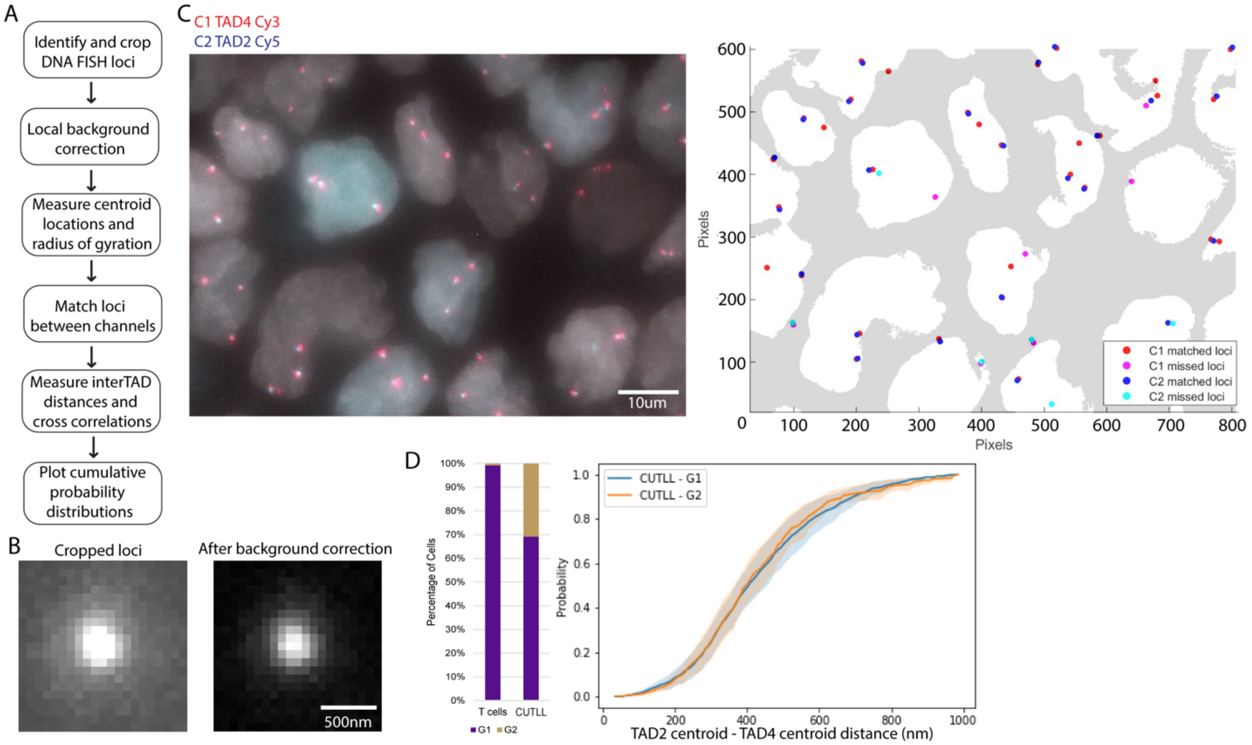


Supplementary Figure 4. Image processing pipeline. Related to Figure 3. A) Overview of the image processing process. First, DNA FISH loci were identified and cropped out from the original image and the background was corrected locally. Second, loci centroid location and radius of gyration are measured. Third, DNA FISH loci from different channels were paired and distances between centroids were measured. Last, the distances were plotted as cumulative probability distributions for each condition. B) Representative images of cropped loci before and after background correction. C) Left: Maximum intensity z-projection of a typical image stack. Right: 2D projection of the locations of matched loci between different channels (C1: TAD4; C2: TAD2; nuclei masks are marked in white). D) Left: bar plot of cells identified in each cell cycle phase in naive T cells and CUTLL1 cells. Right: Cumulative probability distributions of the distance between the centroid of TAD2 and the centroid of TAD4 in CUTLL1 in G1 and G2 phases.

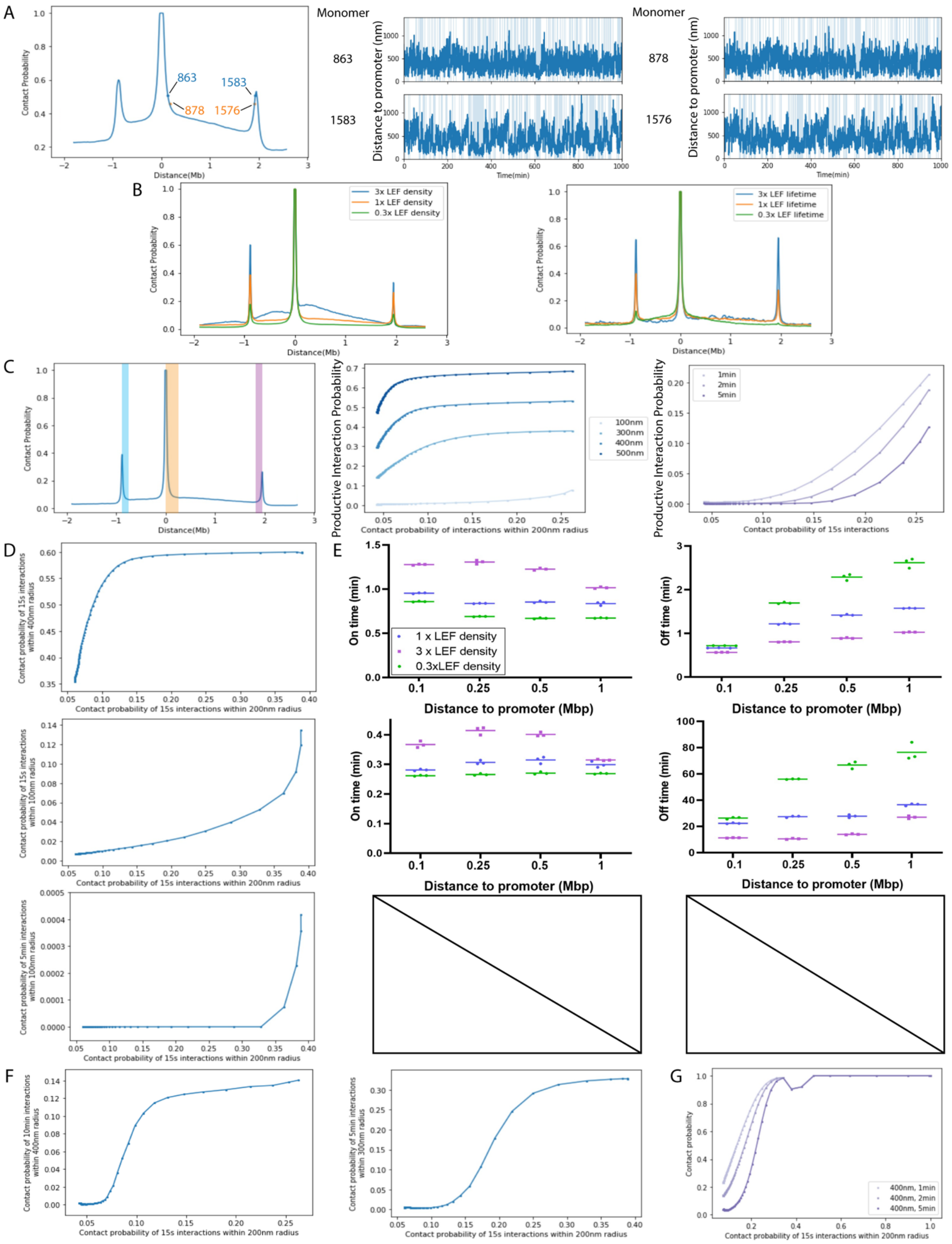
A
Monomer
863
878
1583
1576
Contact Probability
Distance(Mb)
Distance to promoter (nm)
Time(min)
B
3x LEF density
1x LEF density
0.3x LEF density
3x LEF lifetime
1x LEF lifetime
0.3x LEF lifetime
Contact Probability
Distance(Mb)
C
Productive Interaction Probability
Contact probability of interactions within 200nm radius
100nm
300nm
400nm
500nm
1min
2min
5min
Contact probability of 15s interactions
D
Contact probability of 15s interactions within 400nm radius
Contact probability of 15s interactions within 100nm radius
Contact probability of 5min interactions within 100nm radius
Contact probability of 15s interactions within 200nm radius
E
On time (min)
Off time (min)
1 x LEF density
3 x LEF density
0.3xLEF density
Distance to promoter (Mbp)
F
Contact probability of 10min interactions within 400nm radius
Contact probability of 5min interactions within 300nm radius
Contact probability of 15s interactions within 200nm radius
G
Contact probability
400nm, 1min
400nm, 2min
400nm, 5min
Contact probability of 15s interactions within 200nm radius

Supplementary Figure 5. Simulated chromatin dynamics. Related to Figure 4. A) Left: Frequency of contact (<400 nm) with the promoter as a function of genomic position across the locus in simulated data. Two pairs of loci of interest (monomers 863 and 1583, 878 and 1576) exhibit similar contact frequency despite very different distances to the promoter (monomer 811). Center, Right: Example simulated time traces of the distance between loci and the promoter. Pairs of loci (monomers 863 and 1583, 878 and 1576) with similar Hi-C contact frequency are plotted together. B) Frequency of contact (<200 nm) with the promoter (distance origin) as a function of genomic position across the locus in simulated data, in conditions where the LEF density (Left) or LEF lifetime (Right) was changed. C) Left: Frequency of contact (<200 nm) with the promoter as a function of genomic position across the locus in simulated data. Data points from the region marked in cyan plotted in D, F, and Fig. 4, from the region marked in magenta plotted in this panel, and from the region marked in orange plotted in G. Center, Right: The frequency of productive interactions varies nonlinearly as a function of Hi-C contact frequency when imposing a maximum capture radius (Center) or minimum interaction duration (Right). D) Frequency of productive interactions as a function of Hi-C contact frequency when enforcing time- and distance-gating, for other types of possible biologically relevant time- and distance-gatings (productive interaction capture radius and minimum duration for each plot are indicated on the y axis; Hi-C capture radius and minimum duration: 200 nm, 15 s). Top, longer distance-gating only; Middle, shorter distance-gating only; Bottom, shorter distance-gating and longer time-gating. E) Average On- and Off-time durations of productive interactions (capture radius and minimum duration as indicated in D) between the promoter and monomers separated by different genomic distances, for different LEF densities. Each data point is the average of 10 independent simulations. On- and Off-time durations were not plotted for 5 min productive interaction within 100 nm radius due to extremely low contact frequency (< 0.0005). F) Frequency of productive interactions as a function of Hi-C contact frequency when enforcing time- and distance-gating (productive interaction capture radius and minimum duration for each plot are indicated on the y axis; Hi-C capture radius and minimum duration: 200 nm, 15 s). G) Frequency of productive interactions as a function of Hi-C contact frequency when enforcing time- and distance-gating; (productive interaction capture radius and minimum duration for each curve are indicated in the box; Hi-C capture radius and minimum duration: 200 nm, 15 s).

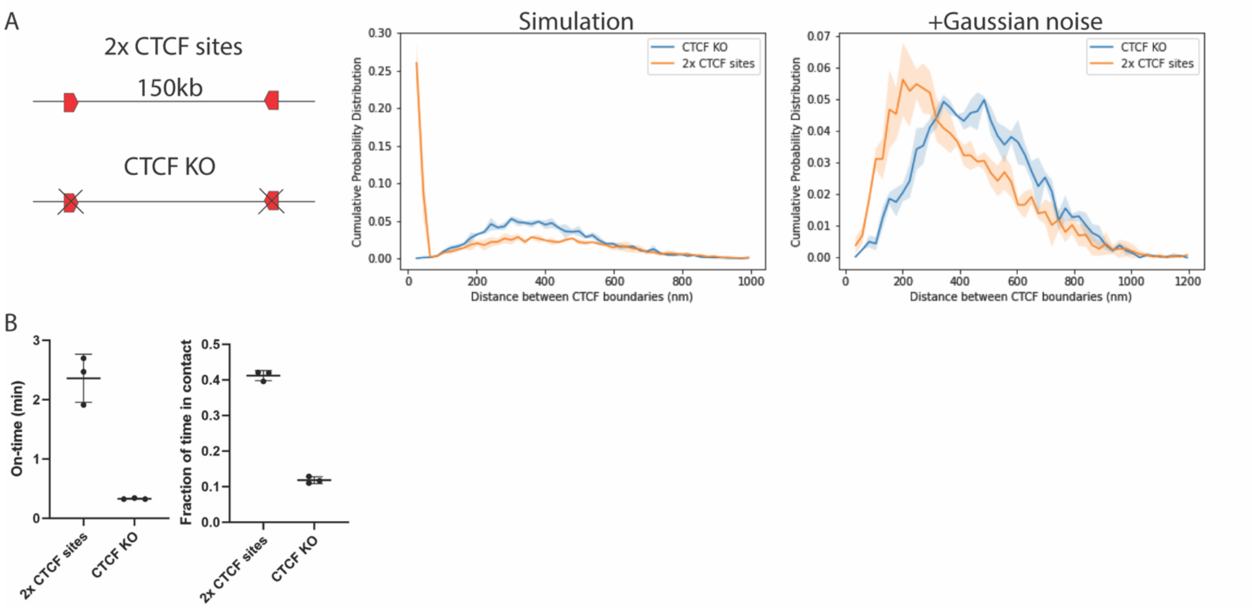


Supplementary Figure 6. Simulations recapitulate dynamics between CTCF boundaries observed in Mach et al (45). A) Distance between two impermeable CTCF binding sites, with and without CTCF. Left, raw simulation data. Right, simulated data with added gaussian noise ($\sigma$ = 94 nm) to simulate localization error in imaging experiments. Shades represent standard deviations of each bin in the cumulative probability distributions (n = 3). B) Simulated on-time duration (left) and fraction of time in contact (right) with and without CTCF.

Supplementary Table 1. DNA FISH probe sequences (see separate document).

Supplementary Table 2. Statistics of cells excluded from cell cycle analysis.

| Cell type | | Total number of cells | Number of cells with more than 4 loci (%) | Number of cells without loci (%) |
|---|---|---|---|---|
| T cells | Sample 1 | 1243 | 4 (0.3%) | 206 (16.6%) |
| | Sample 2 | 247 | 22 (8.9%) | 43 (17.4%) |
| | Sample 3 | 807 | 42 (5.2%) | 566 (70.1%) |
| CUTLL | Sample 1 | 1076 | 27 (2.5%) | 407 (37.8%) |
| | Sample 2 | 1141 | 82 (7.2%) | 519 (45.5%) |

| | | | | |
|---|---|---|---|---|
| | Sample 3 | 814 | 118 (14.5%) | 327 (40.2%) |

Supplementary Table 3. Distance cutoff for interactions used in different publications

| Publication | Distance measurement method | Criterion for interaction | Cell type | Distance cutoff |
|---|---|---|---|---|
| Cardozo Gizzi et al. 2019 | Hi-M | Hi-C | Drosophila | 120 nm |
| Cattoni et al. 2017 | DNA FISH | Hi-C | Drosophila | 120 nm |
| Mateo et al. 2019 | DNA FISH | Hi-C | Drosophila embryo | 150 nm |
| Hafner et al. 2022 | DNA FISH | Hi-C | mESC at E14 stage | 200 nm |
| Murphy and Boettiger, 2022 | DNA FISH | Hi-C | mESC | 150 nm |
| Takei, Zheng, et al. 2021 | seqFISH | Hi-C | Mouse brain cortex | 150 nm (25 kbp resolution)<br>500 nm (1 Mbp resolution) |
| Takei, Yun, et al. 2021 | seqFISH | Hi-C | mESC | 150 nm (25 kbp resolution)<br>500 nm (1 Mbp resolution) |
| Fudenberg and Imakaev, 2017 | Previously published DNA FISH data | Hi-C | GM12878 | 300 nm |
| Wang et al. 2016 | DNA FISH | Hi-C | IMR90 | 500 nm |
| Su et al. 2020 | DNA FISH | Hi-C | IMR90 | 500 nm |
| Chen et al. 2022 | DNA FISH | Hi-C | hESC, human cranial neural crest cells | 250 nm |
| Finn et al. 2019 | DNA FISH | Hi-C | HFF | 350 nm |
